# Functional genomic screens identify human host factors for SARS-CoV-2 and common cold coronaviruses

**DOI:** 10.1101/2020.09.24.312298

**Authors:** Ruofan Wang, Camille R. Simoneau, Jessie Kulsuptrakul, Mehdi Bouhaddou, Katherine Travisano, Jennifer M. Hayashi, Jared Carlson-Stevermer, Jennifer Oki, Kevin Holden, Nevan J. Krogan, Melanie Ott, Andreas S. Puschnik

**Affiliations:** Chan Zuckerberg Biohub, San Francisco, CA, 94158, USA; Gladstone Institutes, San Francisco, CA 94158, USA; University of California San Francisco, Quantitative Biosciences Institute (QBI), San Francisco, CA, 94158, USA; University of California San Francisco, Department of Cellular and Molecular Pharmacology, San Francisco, CA, 94158, USA; Synthego Corporation, Menlo Park, CA 94025

## Abstract

The *Coronaviridae* are a family of viruses that causes disease in humans ranging from mild respiratory infection to potentially lethal acute respiratory distress syndrome. Finding host factors that are common to multiple coronaviruses could facilitate the development of therapies to combat current and future coronavirus pandemics. Here, we conducted parallel genome-wide CRISPR screens in cells infected by SARS-CoV-2 as well as two seasonally circulating common cold coronaviruses, OC43 and 229E. This approach correctly identified the distinct viral entry factors ACE2 (for SARS-CoV-2), aminopeptidase N (for 229E) and glycosaminoglycans (for OC43). Additionally, we discovered phosphatidylinositol phosphate biosynthesis and cholesterol homeostasis as critical host pathways supporting infection by all three coronaviruses. By contrast, the lysosomal protein TMEM106B appeared unique to SARS-CoV-2 infection. Pharmacological inhibition of phosphatidylinositol phosphate biosynthesis and cholesterol homeostasis reduced replication of all three coronaviruses. These findings offer important insights for the understanding of the coronavirus life cycle as well as the potential development of host-directed therapies.

## Introduction

The *Coronaviridae* family includes seven known human pathogens, for which there are no approved vaccines and only limited therapeutic options. The seasonally circulating human coronaviruses (HCoV) OC43, HKU1, 229E and NL63 cause mild, common cold-like, respiratory infections in humans ^1^. However, three highly pathogenic coronaviruses emerged in the last two decades, highlighting the pandemic potential of this viral family ^2–4^. Infection with severe acute respiratory syndrome coronavirus 1 (SARS-CoV-1) and Middle East respiratory syndrome coronavirus (MERS-CoV) can lead to acute respiratory distress syndrome and death, with fatality rates between 10-40% ^5^. SARS-CoV-2, which is currently causing a global pandemic, is less deadly but far more transmissible than SARS-CoV-1 and MERS-CoV, and has been responsible for over 32 million cases and 900,000 deaths globally so far ^5, 6^. Because of the severity of their impact on global health, it is critical to understand how SARS-CoV-2 and other coronaviruses hijack the host cell machinery during infection, and apply this knowledge to develop new therapeutic strategies.

Coronaviruses are enveloped, positive-sense, single-stranded RNA viruses with a genome length of approximately 30kb. Upon receptor binding and membrane fusion, the viral RNA is released into the cytoplasm, where it is translated to produce viral proteins. Subsequently, the viral replication/transcription complexes form on double-membrane vesicles and generate genome copies. These are then packaged into new virions via a budding process, through which they acquire the viral envelope, and the resulting virions are released from infected cells ^7^. During these steps, specific cellular proteins are hijacked and play crucial roles in the viral life cycle. For example, the angiotensin-converting enzyme 2 (ACE2) is exploited as the viral entry receptor for HCoV-NL63, SARS-CoV-1 and SARS-CoV-2 ^8–10^. Additionally, cellular proteases, such as TMPRSS2, cathepsin L and furin, are important for the cleavage of the viral spike (S) protein of several coronaviruses, thereby mediating efficient membrane fusion with host cells ^11–15^. Systematic studies have illuminated virus-host interactions during the later steps of the viral life cycle. For example, proteomics approaches revealed a comprehensive interactome between individual SARS-CoV-2 proteins and cellular proteins ^16, 17^. Additionally, biotin labelling identified candidate host factors based on their proximity to coronavirus replicase complexes ^18^. While these studies uncovered physical relationships between viral and cellular proteins, they do not provide immediate information about the importance of these host components for viral replication.

An orthogonal strategy is to screen for mutations that render host cells resistant to viral infection using CRISPR-based mutagenesis. These screens identify host factors that are functionally required for virus replication and could be targets for host-directed therapies^19^. Several groups have already successfully applied this approach, yet with certain limitations, e.g. the use of Vero green monkey cells instead of a human cell line ^20^, the use of a small CRISPR library only based on the SARS-CoV-2 protein interactome ^21^, or the use of a SARS-CoV-2 harboring a deletion in the spike S1/S2 site due to cell culture adaptation ^22^.

In this study, we have performed a genome-wide CRISPR knockout (KO) screen using wild-type SARS-CoV-2 (USA/WA-1 isolate) in human cells. Importantly, we expanded our functional genomics approach to distantly related *Coronaviridae* in order to probe for commonalities and differences across the family. This strategy can reveal potential pan-coronavirus host factors, thus illuminating targets for antiviral therapy to combat current and potential future outbreaks. We conducted comparative CRISPR screens for SARS-CoV-2 and two seasonally circulating common cold coronaviruses, OC43 and 229E. Our results corroborate previously implicated host pathways, uncover new aspects of virus-host interaction and identify targets for host-directed antiviral treatment.

## Results

### CRISPR knockout screens identify common and virus-specific candidate host factors for coronavirus infection

Phenotypic selection of virus-resistant cells in a pooled CRISPR KO screen is based on survival and growth differences of mutant cells upon virus infection. We chose Huh7.5.1 hepatoma cells as they were uniquely susceptible to all tested coronaviruses. We readily observed drastic cytopathic effect during OC43 and 229E infection (Extended Data Fig. 1a). Huh7.5.1 also supported high levels of SARS-CoV-2 replication but only displayed limited virus-induced cell death (Extended Data Fig. 1b,c). To improve selection conditions for the SARS-CoV-2 CRISPR screen, we overexpressed ACE2 and/or TMPRSS2, which are present at low levels in WT Huh7.5.1 cells (Extended Data Fig. 1d). This led to increased viral uptake of a SARS-CoV-2 spike-pseudotyped lentivirus, confirming the important function of ACE2 and TMPRSS2 for SARS-CoV-2 entry (Extended Data Fig. 1e). We ultimately used Huh7.5.1 cells harboring a bicistronic ACE2-IRES-TMPRSS2 construct for the SARS-CoV-2 screen as these cells sustained efficient infection that led to widespread cell death, while still allowing the survival of a small number of cells (Extended Data Fig. 1c).

The three CRISPR screens - for resistance to SARS-CoV-2, HCoV-229E and HCoV- OC43 - identified a compendium of critical host factors across the human genome (Fig. 1a, Supplementary Table 1). Importantly, the known viral entry receptors ranked among the top hits: ACE2 for SARS-CoV-2 and aminopeptidase N (ANPEP) for 229E (Fig. 1b,c) ^8, 23^. OC43, unlike the other coronaviruses, does not have a known proteinaceous receptor but primarily depends on sialic acid or glycosaminoglycans for cell entry ^24, 25^; consistent with this fact, multiple heparan sulfate biosynthetic genes (B3GALT6, B3GAT3, B4GALT7, EXT1, EXT2, EXTL3, FAM20B, NDST1, SLC35B2, UGDH, XYLT2) were identified in our OC43 screen (Fig. 1d, Extended Data Fig. 2a). Several of these genes were also markedly enriched in the SARS-CoV-2 screen, consistent with a recent report that SARS-CoV-2 requires both ACE2 and cellular heparan sulfate for efficient infection (Fig. 1b, Extended Data Fig. 2a) ^26^. Overall, the identification of the expected entry factors validates the phenotypic selection of our host factor screens.

**Figure 1:**
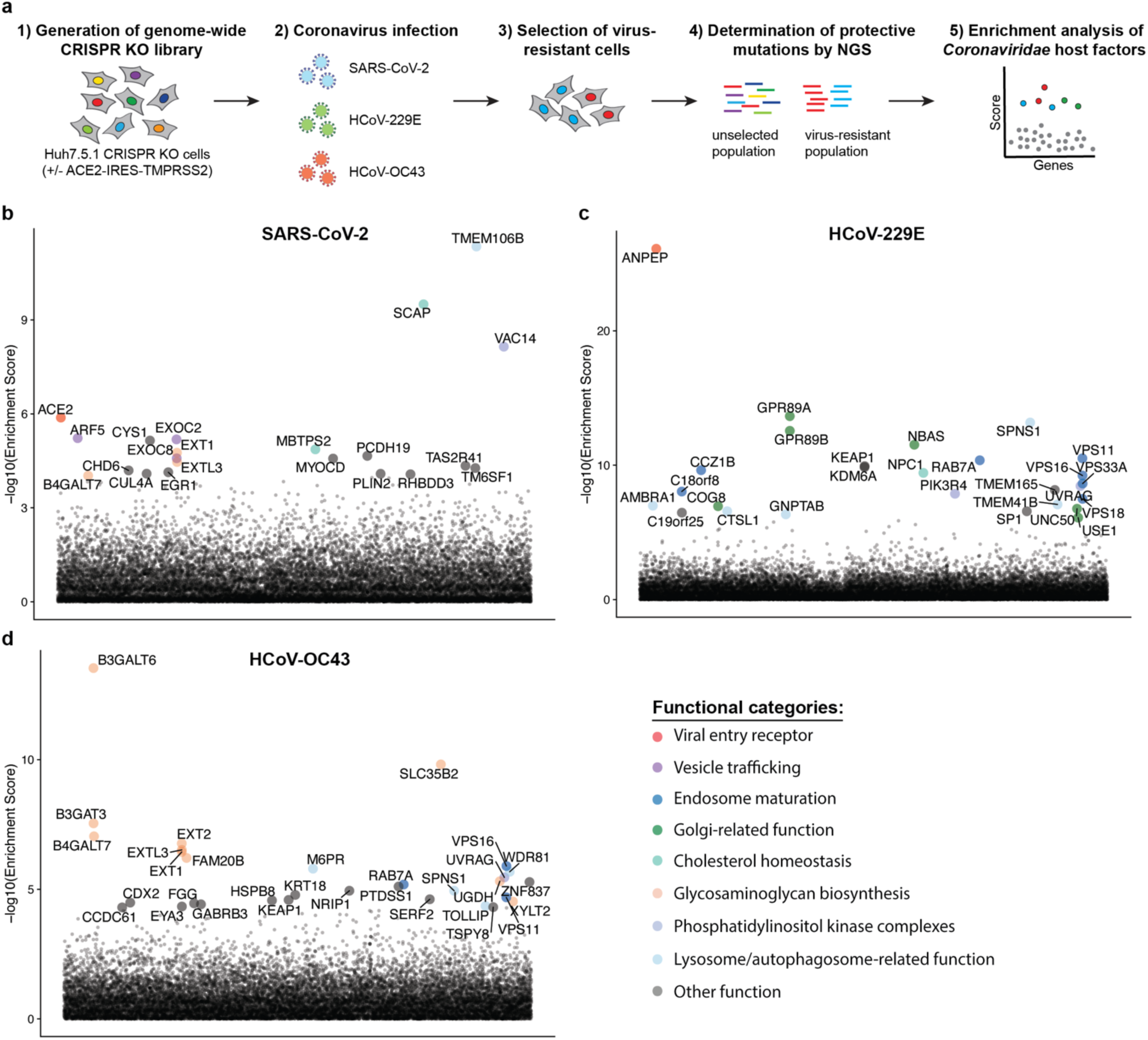
Genome-wide CRISPR KO screens in human cells identify host factors important for infection by for SARS-CoV-2, HCoV-229E and HCoV-OC43. (a) Schematic of CRISPR KO screens for the identification of coronavirus host factors. Huh7.5.1-Cas9 (with bicistronic ACE2-IRES-TMPRSS2 construct for SARS-CoV-2 and without for 229E and OC43 screen) were mutagenized using a genome-wide sgRNA library. Mutant cells were infected with each coronavirus separately and virus-resistant cells were harvested 10-14 days post infection (dpi). The abundance of each sgRNA in the starting and selected population was determined by high-throughput sequencing and a gene enrichment analysis was performed. (b-d) Gene enrichment of CRISPR screens for (b) SARS-CoV-2, (c) 229E and (d) OC43 infection. Enrichment scores were determined by MaGECK analysis and genes were colored by biological function. The SARS-CoV-2 was performed once. The 229E and OC43 screens were performed twice and combined MaGECK scores are displayed.

Gene Ontology (GO) enrichment analysis found a number of cellular processes to be important for multiple coronaviruses. These included proteoglycan and aminoglycan biosynthesis, vacuolar and lysosomal transport, autophagy, Golgi vesicle transport and phosphatidylinositol metabolic processes (Fig. 2a, Supplementary Table 2). In the phosphatidylinositol metabolic process, the SARS-CoV-2 screen identified VAC14, which is part of the PIKfyve kinase complex (Fig. 1b). PIKFYVE itself was moderately enriched in the SARS-CoV-2 screen (Extended Data Fig. 2a). This complex catalyzes the conversion of phosphatidylinositol-3-phosphate to phosphatidylinositol-3,5-bisphosphate, which is localized to late endosomes ^27^. Interestingly, the CRISPR screens with HCoV-229E and HCoV-OC43 identified the subunits (PIK3C3, UVRAG, BECN1 and PIK3R4) of the class III phosphatidylinositol 3-kinase (PI3K) complex, which generates the precursor phosphatidylinositol-3-phosphate in early endosome membranes (Fig. 1c,d and Extended Data Fig. 2a) ^28^. Taken together, our data highlight different steps of the phosphatidylinositol biosynthetic pathway, which regulates endocytic sorting, endomembrane homeostasis and autophagy, to be critical for the life cycle of all three and possibly all coronaviruses.

**Figure 2:**
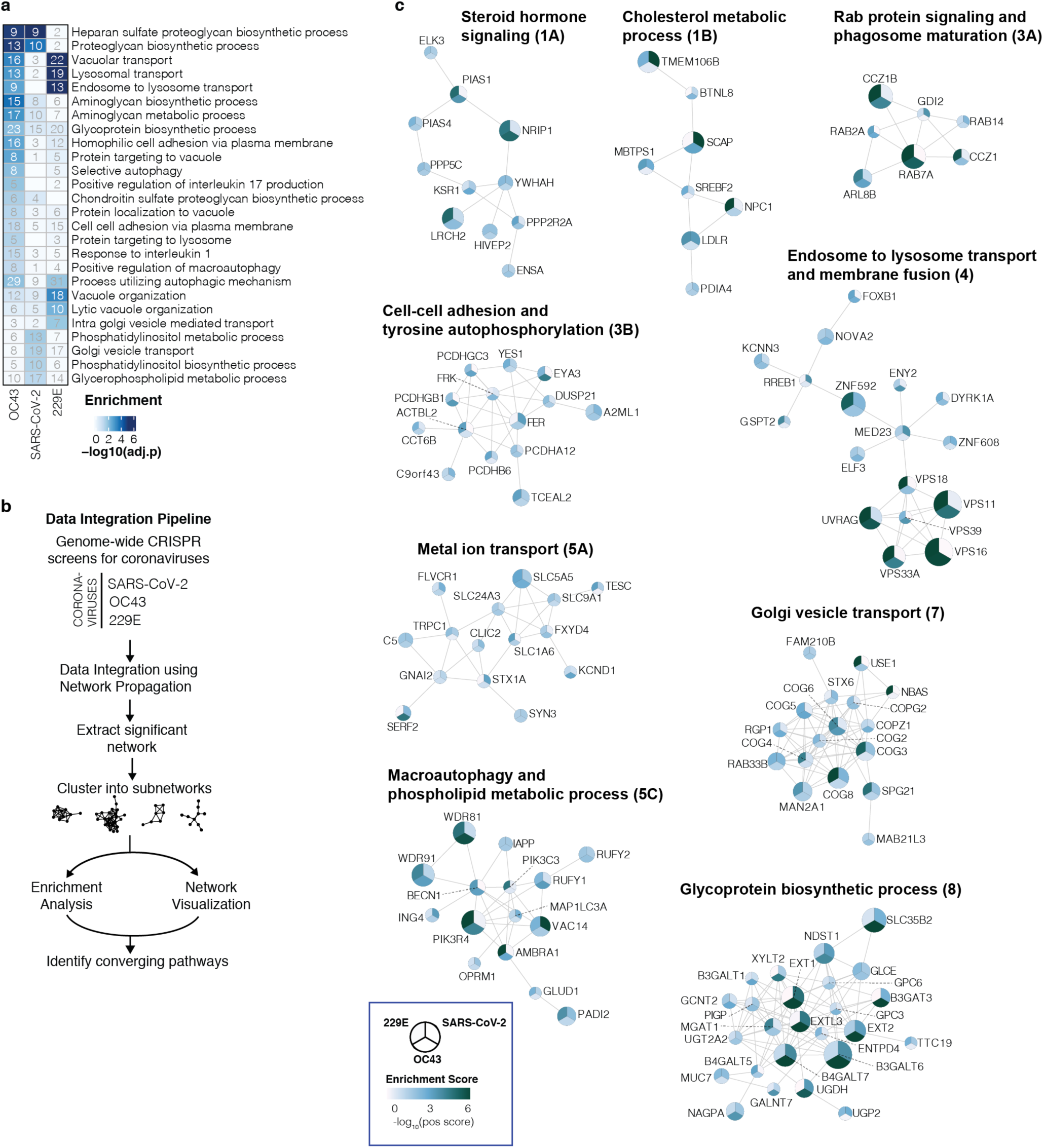
Gene ontology analysis and network propagation highlight pathways and biological signaling networks important for coronavirus infection. (a) Gene ontology (GO) enrichment analysis was performed on significant hits from the individual CRISPR screens (MaGECK enrichment score <= 0.005). P values were calculated by hypergeometric test and a false-discovery rate was used to account for multiple hypothesis testing. The top GO terms of each screen were selected for visualization. **(b)** Data integration pipeline for network propagation of identified host factor genes. Unthresholded positive enrichment scores served as initial gene labels for network propagation using Pathway Commons. Separately propagated networks were integrated gene-wise (via multiplication) to identify biological networks that are shared between all three datasets. Genes found to be significant in the propagation (see Methods) were extracted, clustered into smaller subnetworks, and annotated using GO enrichment analysis. **(c)** Selected biological subnetwork clusters from network propagation. Cluster title indicates the most significant biological function(s) for each cluster. Circle size represents p-value from network propagation permutation test (see Methods). The original positive enrichment score of a gene in each CRISPR screen is indicated by color scale within the circle. The entire set of identified clusters is displayed in Extended Data Fig. 3a. (#) is the cluster number, which refers to the enrichment analysis of biological processes in Extended Data Fig. 3b and Supplementary Table 2.

Another group of genes found in all three CRISPR screens are linked to cholesterol metabolism. The SARS-CoV-2 resistant cell population contained multiple knockouts in genes of the sterol regulatory element-binding protein (SREBP) pathway (SCAP, MBTPS1, MBTPS2) (Fig. 1b, Extended Data Fig. 2a) ^29^. SCAP is an escort protein for the transport of the transcription factors SREBF1 and SREBF2 from the ER to the Golgi in response to low levels of cholesterol. In the Golgi, the SREBF proteins are sequentially cleaved by the proteases MBTPS1 and MBTPS2. Subsequently, the transcription factors translocate to the nucleus to activate fatty acid and cholesterol biosynthesis. SREBF1 and SREBF2 themselves did not score in the screen, potentially due to their functional redundancy. LDLR (Low Density Lipoprotein Receptor), important for cholesterol uptake, was also enriched in both the SARS-CoV-2 and the 229E screen, while SCAP was enriched in the OC43 screen (Extended Data Fig. 2a,b). Additionally, NPC1 (Niemann– Pick intracellular cholesterol transporter 1), which facilitates export of cholesterol from the endolysosomal compartment, ranked highly in the 229E screen (Fig. 1c) ^30^. Overall, our data indicates a strong link between intracellular cholesterol levels and infection by all three coronavirus.

Some genes were found in the OC43 and 229E screens, but not in the SARS-CoV-2 screen. For instance, the common cold coronavirus screens showed a strong overlap of genes, which are important for endosome and autophagosome maturation (Fig. 1c,d and Extended Data Fig. 2b). These include the small GTPase Rab7a, components of the HOPS complex (VPS11, VPS16, VPS18, VPS33A), the Ccz1-Mon1 guanosine exchange complex (CCZ1, CCZ1B, C18orf8), the WDR81-WDR91 complex, and other genes related to lysosome and autophagosome function (SPNS1, TOLLIP, TMEM41B, AMBRA1) ^31–37^. We also identified cathepsin L (CTSL1) as well as the mannose-6-phosphate receptor (M6PR) and GNPTAB, which are important for proper trafficking of lysosomal enzymes from the trans-Golgi network ^38, 39^. Interestingly, the HOPS complex, cathepsins, GNPTAB and SPNS1 were previously linked to Ebola virus entry, implying similar viral entry strategies ^39, 40^.

The absence of endolysosomal factors in the SARS-CoV-2 screen may be explained by the ectopic expression of the cell-surface protease TMPRSS2 in this screen. The cleavage of SARS-CoV-2 spike can occur either at the plasma membrane via TMPRSS2 or in endolysosomes through cathepsins. Sufficient TMPRSS2 levels may thus ablate the requirement for cathepsin and other factors linked to endolysosomal activity ^11^. Consistent with this hypothesis, a CRISPR screen using a SARS-CoV-2 with a spike S1/S2 site deletion, which preferentially uses the endolysosomal entry route, showed strong enrichment in RAB7A, the Ccz1-Mon1 guanosine exchange complex, the HOPS complex and the WDR81-WDR91 complex, similar to the OC43 and 229E screen results ^22^. This suggests that endolysosomal host factors can be required for infection under certain conditions but may become largely dispensable for SARS-CoV-2, especially in TMPRSS2-expressing cell types. As nasal and lung epithelial cells can express high levels of TMPRSS2 ^41^, we speculate that the genes identified in the SARS-CoV-2 CRISPR screen using Huh7.5.1-ACE2-IRES-TMPRSS2 cells may represent rate-limiting factors that are more physiologically relevant to SARS-CoV-2 infection *in vivo* than the endolysosomal components found in our other two screens and in other studies.

The OC43 and 229E screens also uncovered KEAP1, the principal negative regulator of NRF2, whose activation restores cellular redox and protein homeostasis (Fig. 1c,d) ^42^. Activation of the NRF2 transcriptional program may induce a cellular state that is protective against coronavirus infection. Indeed, NRF2 agonists seem to elicit an antiviral response as demonstrated in cell culture and were proposed for SARS-CoV-2 treatment^43, 44^.

In addition to genes that scored in multiple CRISPR screens, we also found genes that were only enriched in one screen. Several genes related to the Golgi apparatus were uncovered only in the 229E screen and may possibly have 229E-specific roles. Among them were GPR89A and GPR89B, which encode two highly homologous G protein coupled receptors important for Golgi acidification ^45^, and NBAS and USE1, which play a role in Golgi-to-ER retrograde transport ^46^. The exact role of these factors in coronavirus infection – and their specificity to 229E – remain to be determined.

The SARS-CoV-2 screen identified multiple subunits of the exocyst (EXOC1-8) (Fig. 1b, Extended Data Fig. 2a), an octameric protein complex that facilitates the tethering of secretory vesicles to the plasma membrane prior to SNARE-mediated fusion ^47^. This complex could therefore facilitate trafficking of virus particles during entry or egress. The top hit of the SARS-CoV-2 screen was TMEM106B, a poorly characterized lysosomal transmembrane protein linked to frontotemporal dementia (Fig. 1b) ^48^. Deletions in TMEM106B caused defects in lysosome trafficking, impaired acidification and reduced levels of lysosomal enzymes but its precise molecular function remains enigmatic ^48, 49^. TMEM106B KO could thus indirectly affect SARS-CoV-2 entry, although it is also possible that SARS-CoV-2 physically interacts with TMEM106B, for example as lysosomal receptor, similar to NPC1 and LAMP1 for Ebola and Lassa virus, respectively ^40, 50^.

Overall, the comparative CRISPR screen strategy provides a rich list of shared and distinct candidate host factors for subsequent validation and host-directed inhibition of coronavirus infection.

### Network propagation across multiple CRISPR screens highlights functional biological clusters important for coronavirus infection

To better understand the functional connections between the genes identified in our screens, we performed network propagation (Fig. 2b) ^51^. This approach integrates large datasets using networks, thereby identifying subnetworks and pathways that are common across datasets. Propagations from the three CRISPR screens identified subnetworks most common to all three viruses and independently confirmed the biological processes highlighted as important for coronavirus infection in our previous analysis (Fig. 2c, Extended Data Fig. 3, Supplementary Table 2 and 3). For instance, we found clusters linked to cholesterol metabolism (containing SCAP, MBTPS1, SREBF2, LDLR and NPC1), endosome to lysosome transport (including the HOPS complex components VPS11, VPS16, VPS18, VPS33A and VPS39) and glycoprotein biosynthetic processes (containing heparan sulfate biosynthesis genes). Another cluster reflected the critical role of autophagy/ phospholipid metabolism and indicated a functional link between VAC14 and subunits of the PI3K complex as described above.

Moreover, network propagation also identified previously unappreciated biological functions, such as steroid hormone signaling, cell-cell adhesion, metal ion transport, intra-Golgi vesicle transport, snare complex assembly, Rab protein signal transduction, peroxisomal transport and mRNA splicing (Fig. 2c, Extended Data Fig. 3, Supplementary Table 2 and 3). Altogether, network propagation revealed numerous distinct cellular processes that may have critical roles during coronavirus infection.

### Knockout of candidate host factor genes reduces coronavirus replication

To validate the candidate genes from the SARS-CoV-2 screen, we generated individual KO cells in two cell types. We first introduced gene deletions for several top hits in A549 lung epithelial cells transduced with ACE2 (A549-ACE2) using Cas9 ribonucleoproteins (RNPs) (Extended Data Fig. 4a). SARS-CoV-2 RNA levels were markedly reduced in cells that contained indel mutations in ACE2, the ADP Ribosylation Factor 5 (ARF5), multiple subunits of the exocyst (EXOC2/6/8), the cholesterol homeostasis genes SCAP, MBTPS1 and MBTPS2, the phosphatidylinositol kinase complex components PIKFYVE and VAC14, as well as TMEM106B (Fig. 3a). Additionally, we generated clonal Huh7.5.1 cells (without the ACE2-IRES-TMPRSS2 construct) harboring frameshift mutations in a subset of the same genes (Extended Data Fig. 4b). Deletion of TMEM106B and VAC14 decreased SARS-CoV-2 replication, and this effect was reversed by cDNA complementation (Fig. 3b,c), thus confirming the role of these two factors in the SARS-CoV-2 life cycle. Similarly, knocking out SCAP, MBTPS2 or EXOC2 led to a decrease of SARS-CoV-2 RNA levels (Fig. 3d). When we infected the same Huh7.5.1 KO cells with HCoV-OC43 and HCoV-229E, we observed reduced viral replication in SCAP and MBTPS2 KO cells, but not in TMEM106B KO and only moderately in VAC14 KO cells (Fig. 3e). This suggests that the latter genes are more rate-limiting in SARS-CoV-2 infection.

**Figure 3:**
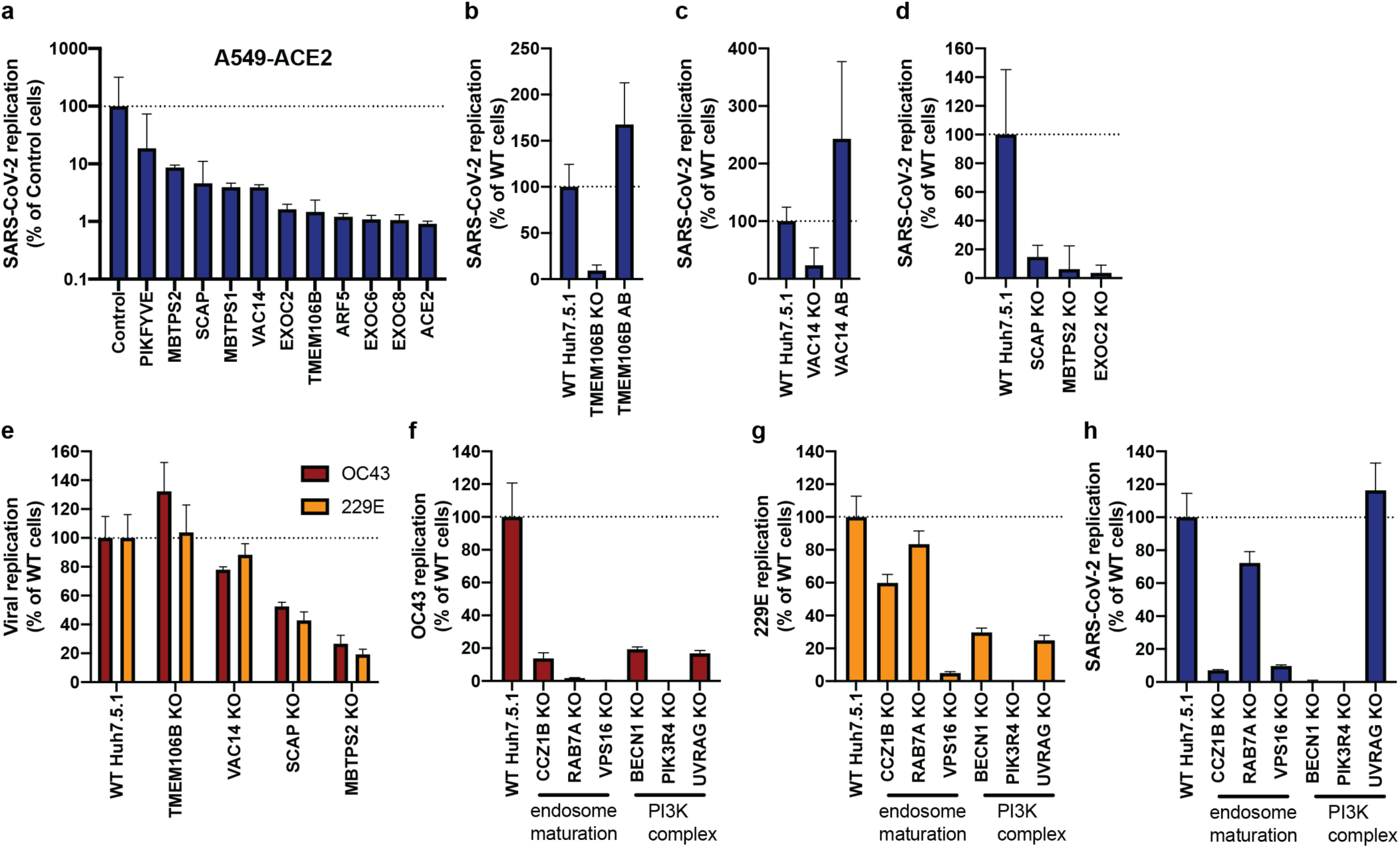
Knockout of candidate host factor genes reduces coronavirus infection. **(a)** RT-qPCR quantification of intracellular SARS-CoV-2 levels in RNP edited A549-ACE2 cells. Cells were infected using moi=0.01 and harvested at 72 hours post infection (hpi). **(b-d)** RT-qPCR quantification of intracellular SARS-CoV-2 levels in WT Huh7.5.1 cells or cells harboring frameshift mutations or frameshift mutant cells complemented with respective cDNAs. Cells were infected using moi=0.01 and harvested at 24 hpi. **(e-g)** RT-qPCR quantification of intracellular OC43 and 229E RNA levels in WT and KO Huh7.5.1 cells. Cells were infected using moi=0.05 (229E) and moi=3 (OC43) and harvested at 48 hpi. **(h)** RT-qPCR quantification of intracellular SARS-CoV-2 levels in Huh7.5.1 WT or KO cells by RT-qPCR. Cells were infected using moi=0.01 and harvested at 24 hpi. For SARS-CoV-2 infection, viral transcripts were normalized to cellular RNaseP. For OC43 and 229E experiments, viral RNA was normalized to 18S RNA. For all RT-qPCR experiments, results are displayed relative to infection in WT cells and data represent means ± s.e.m. from 3 biological samples.

Finally, we probed cells lacking several genes involved in endosome maturation or the PI3K complex, which were initially found in the common cold coronavirus screens. We saw reduced viral replication for OC43 and 229E (Fig. 3f,g). Additionally, we observed increased cell viability in all KO cells relative to WT Huh7.5.1 cells 8 dpi, indicating that endosome maturation is important for infection by the common cold viruses and for virus-induced cell death (Extended Data Fig. 4c). Next, we tested whether the hits shared between OC43 and 229E might also affect SARS-CoV-2. Indeed, SARS-CoV-2 infection also depended on endosomal factors in the context of Huh7.5.1 without ACE2-IRES-TMPRSS2, similar to the common cold coronaviruses (Fig. 3h). Additionally, deletions of the PI3K subunits, in particular PIK3R4 KO, led to a strong decrease in replication of all three coronaviruses (Fig. 3h). Together, these experiments confirm that the host factors identified in our screens have functional roles for *Coronaviridae* and demonstrate that important aspects of SARS-CoV-2 biology can be revealed by studying the common cold coronaviruses.

### Compounds directed at host factors inhibit coronavirus replication

Host factors important for virus infection are potential targets for antiviral therapy. Host-directed therapy is advantageous as it allows pre-existing drugs to be repurposed, it may provide broad-spectrum inhibition against multiple viruses, and it is generally thought to be more refractory to viral escape mutations than drugs targeting viral factors ^52^. We therefore explored whether the cellular pathways identified in our screens could serve as targets for therapy against coronavirus infection.

Given the strong dependence of all three coronaviruses on PIK3R4, we tested SAR405, a selective and ATP-competitive inhibitor of class III PI3K (PIK3C3) ^53^. The drug exhibited a dose-dependent effect against all three coronaviruses with low cytotoxicity, consistent with the reduced virus replication in PIK3R4 KO cells, and suggesting that it could serve as a pan-coronavirus inhibitor (Fig. 4a). As VAC14, a PIKfyve complex component, was a strong hit in the SARS-CoV-2 screen, we also tested the PIKfyve inhibitor YM201636 and observed inhibition of SARS-CoV-2 replication (Fig. 4b) ^54^. Similar antiviral activity was previously demonstrated with apilimod, another PIKfyve inhibitor ^55–57^.

**Figure 4:**
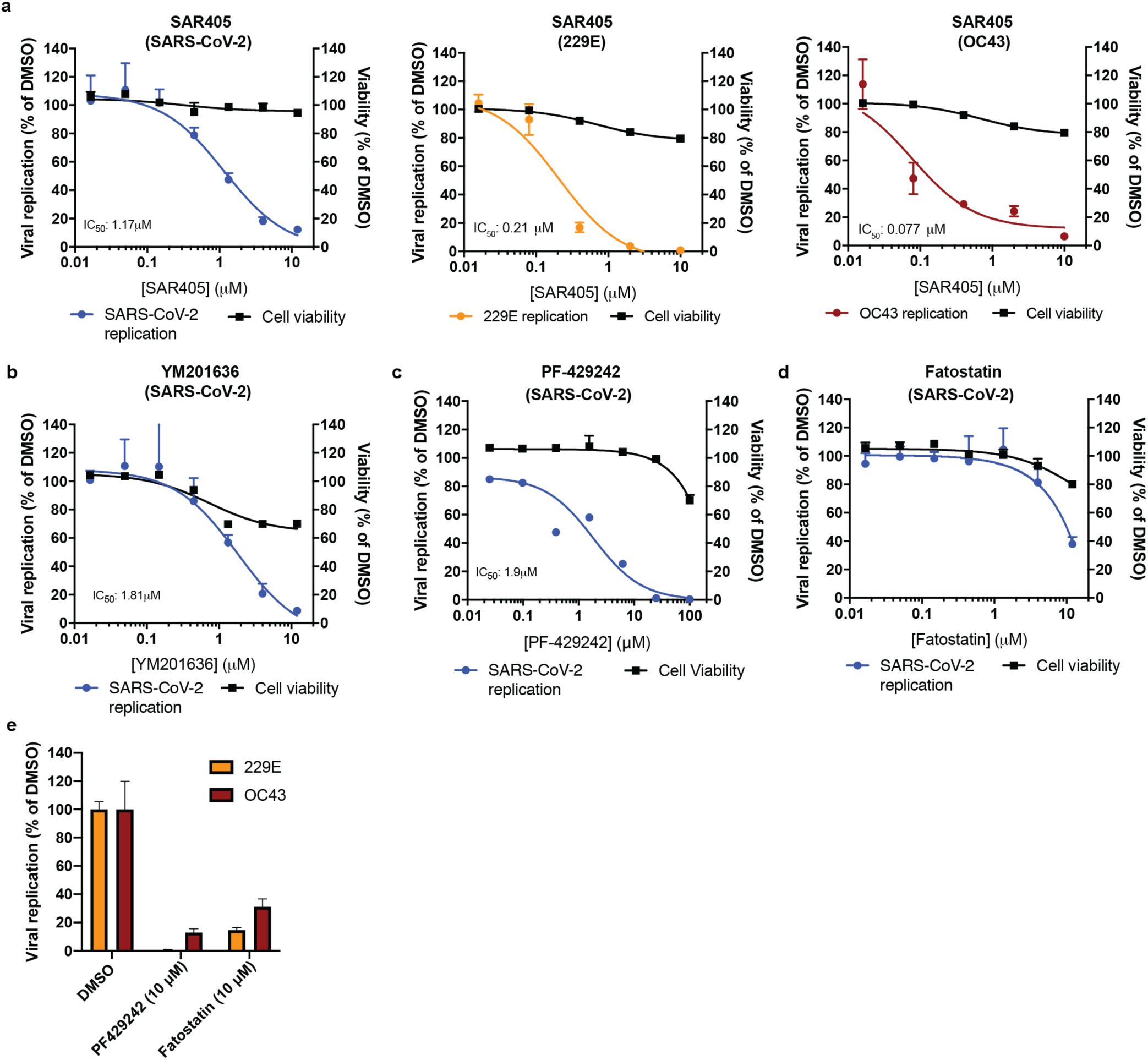
Pharmacological inhibition of phosphatidylinositol kinase complexes and cellular cholesterol homeostasis decreases infection with SARS-CoV-2 and common cold coronaviruses. **(a)** SAR405 dose-response curves for SARS-CoV-2, HCoV-229E and HCoV-OC43 replication in Huh7.5.1 cells and for cell viability of drug treated cells. **(b-d)** Dose-response curves of the effect of **(b)** YM201636, **(c)** PF-429242, and **(d)** fatostatin to on SARS-CoV-2 replication in Huh7.5.1 cells and on cell viability of drug treated cells. **(e)** Quantification of 229E and OC43 replication in the presence of PF-429242 or fatostatin. For all experiments, compounds were added simultaneously with virus. Viral RNA was quantified after 24 hpi (SARS-CoV-2) or 48hpi (229E and OC43) using RT-qPCR and normalized to RnaseP (SARS-CoV-2) or 18S RNA (229E and OC43). Values represent means ± s.e.m. relative to DMSO treated cells. For cell viability, datasets represent means ± s.d. and values are relative to DMSO treated uninfected controls. Non-linear curves were fitted with least squares regression using GraphPad Prism 8 and IC_50_ was determined for (a-c). All experiments were performed in 3 biological replicates.

Furthermore, we tested compounds modulating cholesterol homeostasis as this pathway appeared important for all three coronaviruses as well. PF-429242, a reversible, competitive aminopyrrolidineamide inhibitor of MBTPS1 showed dose-dependent reduction of SARS-CoV-2 replication with cytotoxicity only at high concentration (Fig. 4c)^58^. Fatostatin, which binds to SCAP and inhibits ER-to-Golgi translocation of SREBPs, also moderately reduced SARS-CoV-2 infection levels at higher doses (Fig. 4d) ^59^. The two cholesterol modulating compounds also led to a decrease in OC43 and 229E levels, suggesting that modulation of intracellular cholesterol levels could be used as pan-coronavirus treatment (Fig. 4e). Therefore, our genetic and pharmacological data suggest that both phosphatidylinositol kinase complexes and cholesterol homeostasis are potential targets for pan-coronavirus host-directed therapy in vitro and may be explored further in vivo.

## Discussion

In this study, we performed genome-scale CRISPR KO screens to identify host factors important for SARS-CoV-2, HCoV-229E and HCoV-OC43. Our data highlight that while the three coronaviruses exploit distinct entry factors, they also depend on a convergent set of host pathways, with potential roles for the entire *Coronaviridae* family. In particular, genes involved in cholesterol homeostasis were enriched in all screens and in the network propagation. Consistent with our data, two SARS-CoV-2 interactomes revealed binding of viral proteins to SCAP, and a recent CRISPR screen focused on the interactome components also identified SCAP as a host factor critical for SARS-CoV-2 replication ^16, 17, 21^; given the essentiality of SCAP for replication, viral proteins are likely to positively regulate SCAP activity and cholesterol levels. Cellular cholesterol homeostasis has previously been linked to viral entry and membrane fusion in the context of bunya- and hantavirus infections, suggesting a pro-viral function across different viral families ^60–62^. Consistent with this hypothesis, treatment with 25-hydroxycholesterol, which blocks SREBP processing and stops cholesterol synthesis, reduced infection with SARS-CoV-1 and CoV-2 spike-pseudotyped viruses ^29, 63^.

Additionally, phosphatidylinositol biosynthesis was uncovered as an important pathway for coronavirus infection. While PIKfyve has previously been implicated through chemical inhibition ^55–57^, we identified the upstream PI3K complex as a new critical host factor, potentially exhibiting pan-coronavirus function. Due to its involvement in multiple cellular processes including vesicular trafficking and autophagy ^28^, it remains to be determined whether coronaviruses hijack this pathway during entry or for the generation of double-membrane vesicles required for the viral replication/transcription complexes.

Our results also inform those of a recent drug repurposing screen, which identified ∼100 compounds that inhibited SARS-CoV-2 replication ^64^; notably, among those were PIKfyve inhibitors, protease inhibitors and modulators of cholesterol homeostasis. Our functional genomics data therefore suggest that the observed effects of several compounds were possibly due to inhibition of critical host factors.

In conclusion, our study offers important insight into host pathways commonly hijacked by coronaviruses. Importantly, the identification of the phosphatidylinositol PIK3C3 kinase complex as a potent therapeutic target for SARS-CoV-2 based on the 229E and OC43 screens underscores the value of the parallel CRISPR screening approach for finding novel therapies against SARS-CoV-2 and other *Coronaviridae*.

---

**Extended Data Figure 1:**
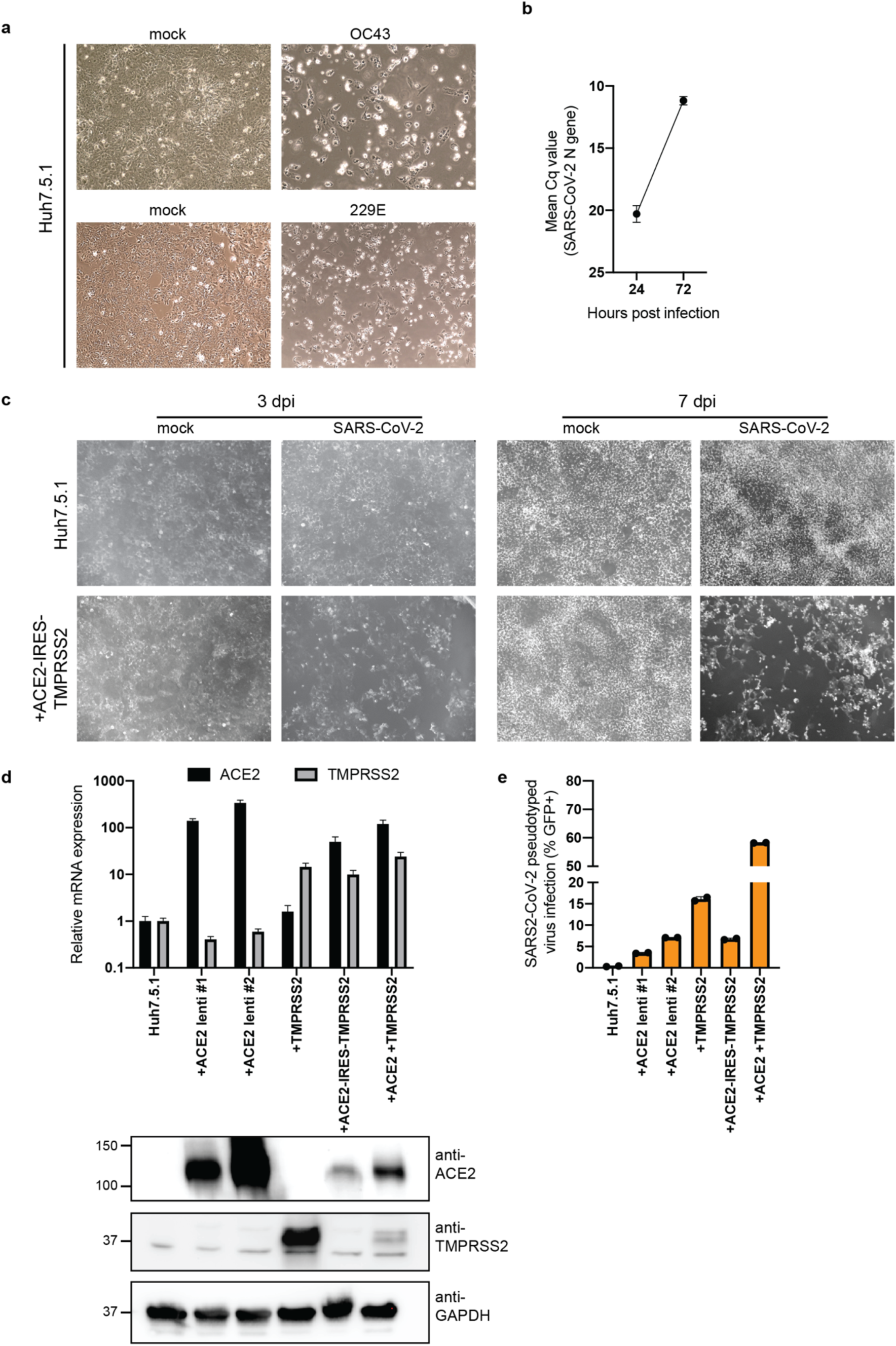
Huh7.5.1 cells are susceptible to SARS-CoV-2, HCoV-OC43 and HCoV-229E. **(a)** Light microscopy images of WT Huh7.5.1 infected with OC43 (7 dpi) and 229E (4 dpi). **(b)** Quantification of SARS-CoV-2 RNA in WT Huh7.5.1 cells at 24 and 72 hpi by RT-qPCR. Cq values represent mean ± s.e.m. from 3 biological replicates. **(c)** Light microscopy images of SARS-CoV-2 infected WT Huh7.5.1 cells or Huh7.5.1 cells expressing ACE2-IRES-TMPRSS2 at 3 and 7 dpi. **(d)** Quantification of ACE2 and TMPRSS2 expression in WT and lentivirally transduced Huh7.5.1 cells by RT-qPCR and Western blot. mRNA levels are displayed as mean ± s.e.m. from two independent harvests and are relative to expression in WT cells. Anti-ACE2 and anti-TMPRSS2 antibodies were used to detect protein levels in WT and overexpression cells. GAPDH was used as loading control. Molecular weight markers are indicated on the left. **(e)** Quantification of infection with pseudotyped lentivirus bearing SARS-CoV-2 spike and expressing a GFP by flow cytometry. Values are from two biological samples and are displayed as means ± s.d.

**Extended Data Figure 2:**
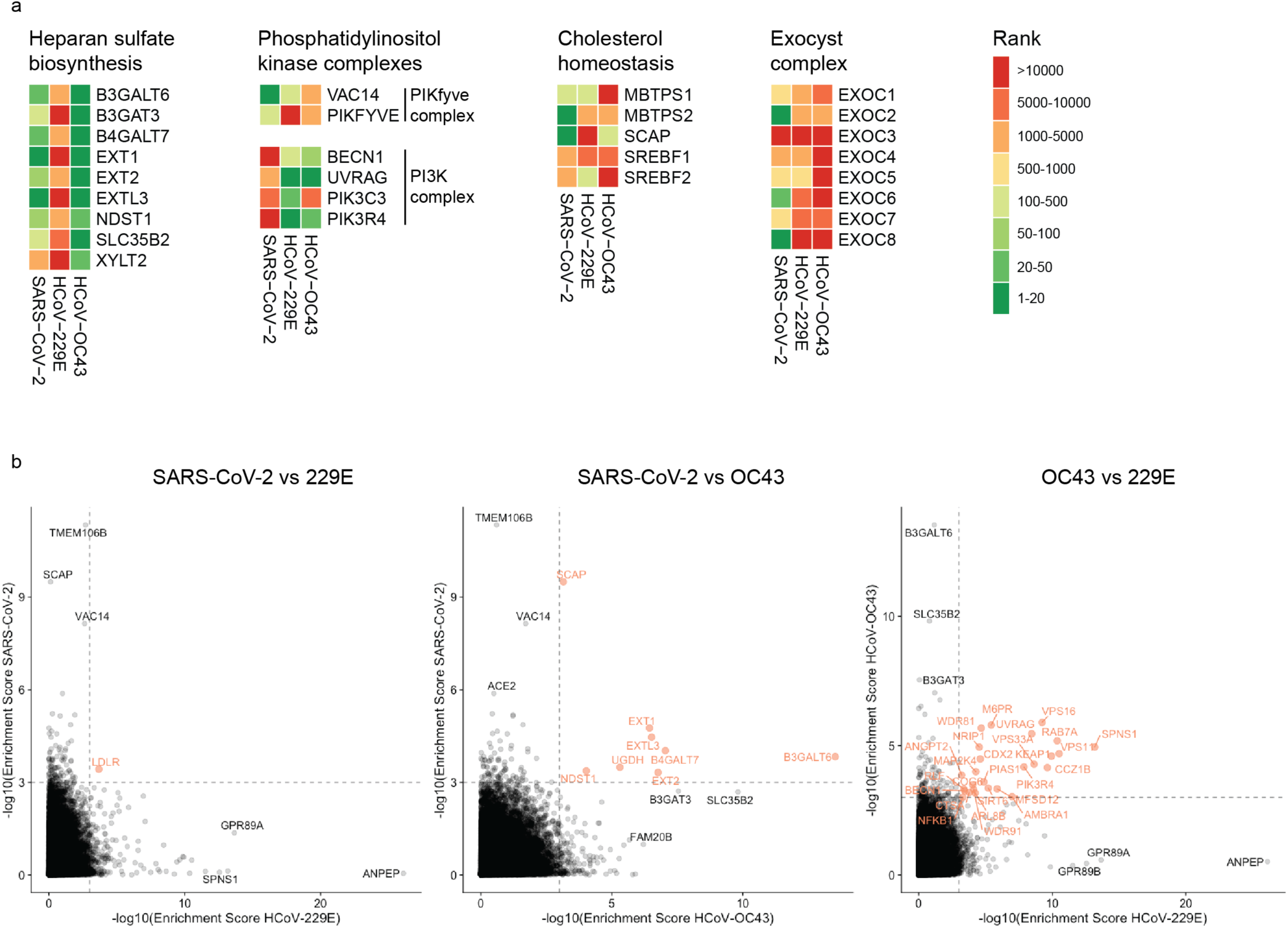
Comparison of CRISPR screens reveals common and distinct host factors across SARS-CoV-2, 229E and OC43. **(a)** CRISPR screen ranking of genes, which are part of specific cellular pathway or complexes, across the three CRISPR screens. **(b)** Pairwise comparisons of gene enrichments between two CRISPR screens. Dotted lines indicate -log_10_(Enrichment score) > 3. Genes that scored above the threshold in both screens, are highlighted in red.

**Extended Data Figure 3:**
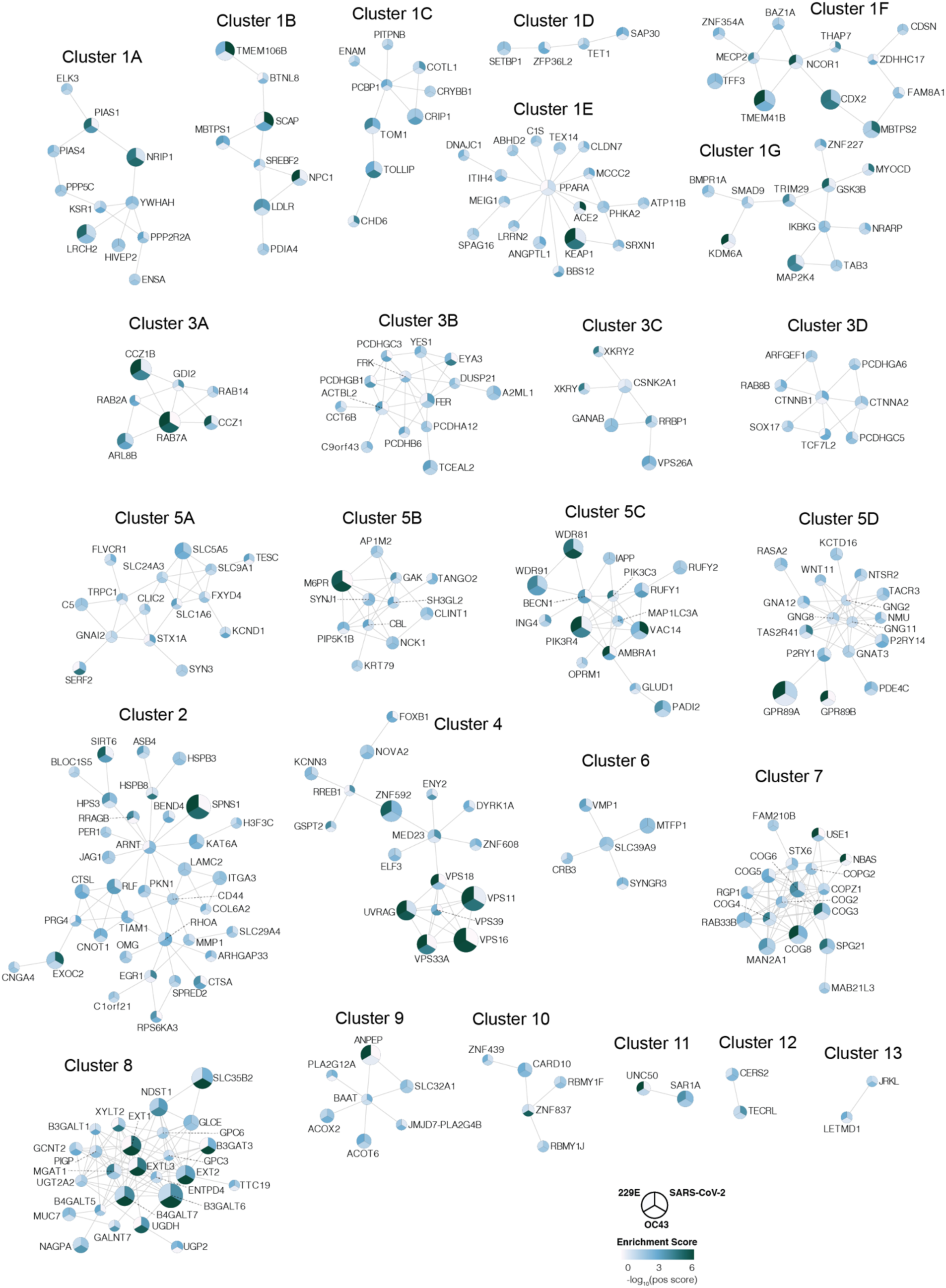
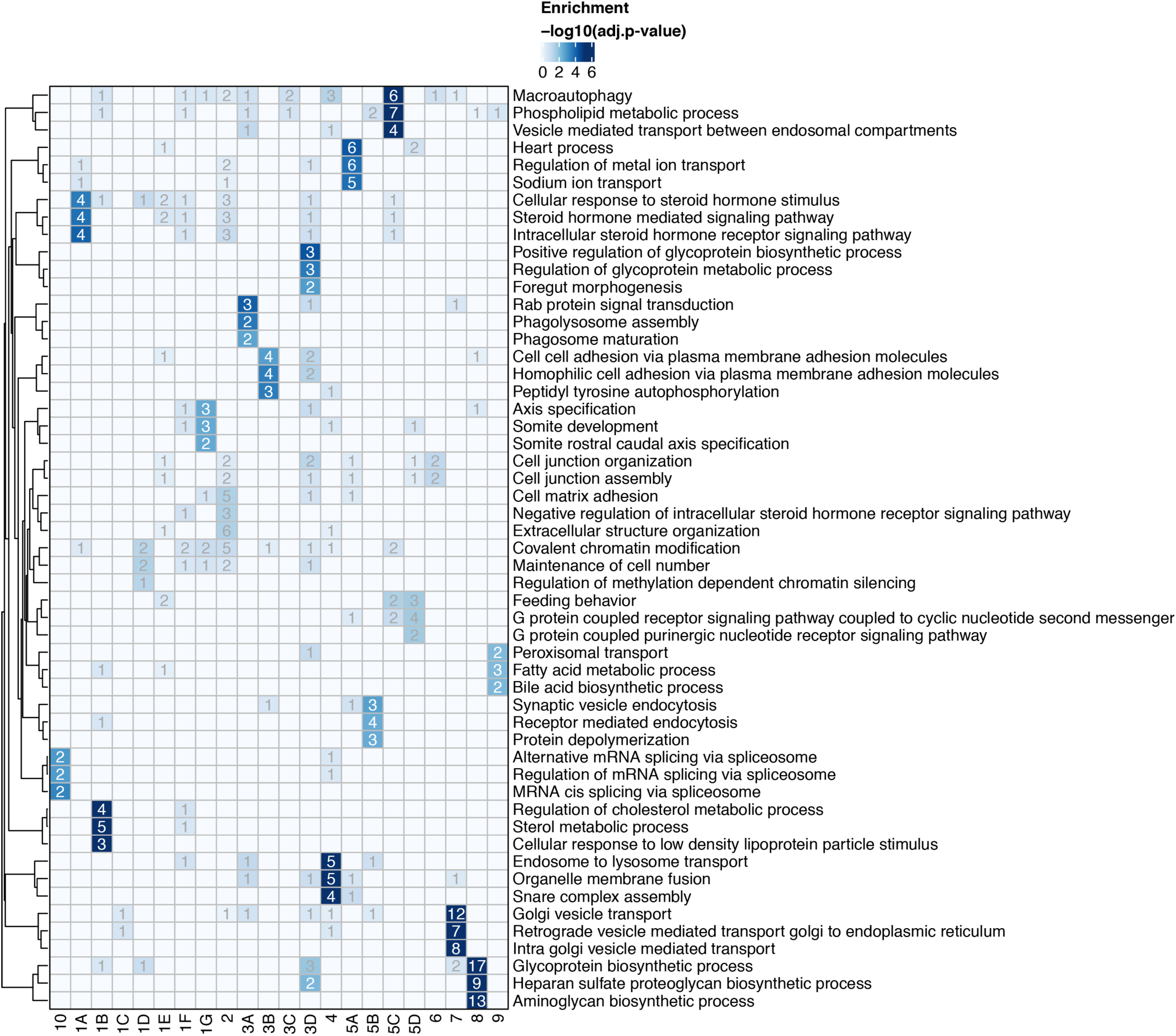
Network propagation of CRISPR screen hits reveals functional clusters with distinct biological functions. **(a)** Biological subclusters from network propagation. Cluster number refers to the enrichment analysis of biological processes for each cluster, displayed in Extended Data Fig. 3b. Circle size represents p-value from integrative network propagation permutation test (gene-wise multiplication across datasets, see Methods). The original positive enrichment score of a gene in each CRISPR screen is indicated by color scale within the circle. **(b)** Gene ontology (GO) enrichment analysis was performed on each subcluster from the network propagation. P values were calculated by hypergeometric test and a false-discovery rate was used to account for multiple hypothesis testing. The entire set of enriched biological processes for each subcluster is listed in Supplementary Table 2.

**Extended Data Figure 4:**
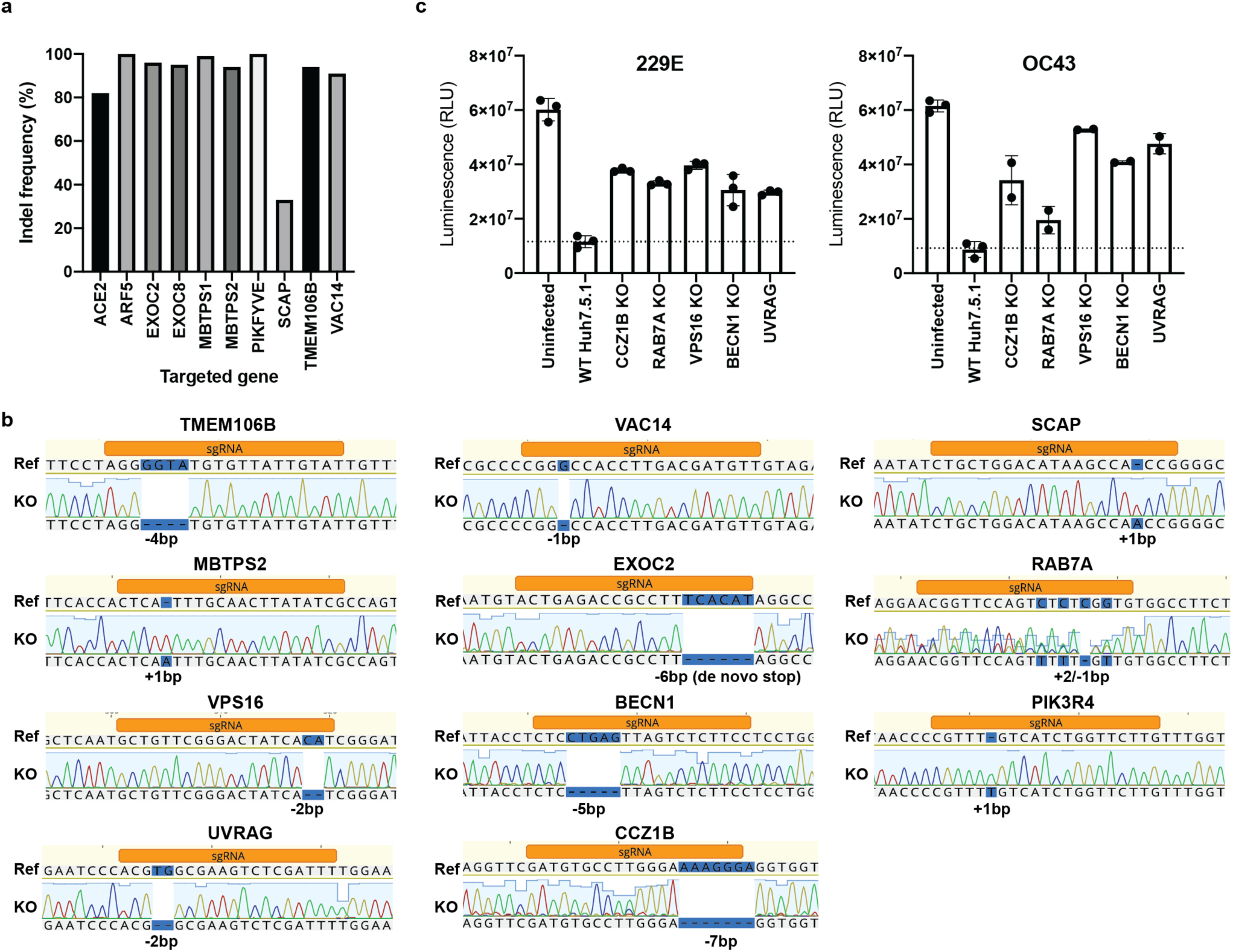
Knockout of host factor genes reduces coronavirus infection and virus-induced cell death. **(a)** Indel frequency of RNP-edited polyclonal A549-ACE2 KO cells. Targeted loci were PCR-amplified, Sanger-sequenced and analyzed using Inference of CRISPR Edits (ICE) analysis ^65^. **(b)** Genotyping of clonal Huh7.5.1. Targeted loci were PCR-amplified, Sanger-sequenced and aligned to WT reference sequence. Frameshifts are highlighted in blue. **(c)** Cell viability measurement of 229E or OC43 infected WT and KO Huh7.5.1 cells. Cells were infected with 229E (moi=0.05) or OC43 (moi=3) and viability was determined 8 dpi using Cell Titer Glo. Values are displayed as means ± s.d. from three (229E) or two (OC43) biological samples.

## Material and Methods

### Cell culture

Huh7.5.1 (gift from Frank Chisari), HEK293FT (Thermo Scientific), Vero cells (ATCC) and A549-ACE2 cells (gift from Olivier Schwartz) were cultured in DMEM (Gibco) supplemented with 10% fetal bovine serum (FBS, Omega Scientific), penicillin/streptomycin (Gibco), non-essential amino acids (Gibco) and L-glutamine (Gibco) at 37C and 5% CO_2_. Cell lines were tested negative for mycoplasma contamination.

### Plasmids, cloning and lentivirus production

The following cDNA sequence containing plasmids were obtained: hACE2 (Addgene, #1786, gift from Hyeryun Choe), TMPRSS2 (Addgene, #53887, gift from Roger Reeves), TMEM106B (Genscript, OHu17671) and VAC14 (Addgene, #47418, gift from Peter McPherson).

Individual cDNAs were cloned into EcoRV-cut plenti-CMV-Puro-DEST Addgene, #17452, gift from Eric Campeau & Paul Kaufman) or plenti-CMV-Hygro-DEST (Addgene, #17454, gift from Eric Campeau & Paul Kaufman) using NEBuilder HiFi DNA Assembly Master Mix (NEB). To generate the plenti-CMV-ACE2-IRES-TMPRSS2 construct, ACE2, EMCV IRES (derived from pLenti-DsRed_IRES_EGFP (Addgene, #92194, gift from Huda Zoghbi)), and TMPRSS2 were individually amplified with addition of overlapping sequences and the three fragments were assembled using NEBuilder HiFi DNA Assembly Master Mix.

Lentivirus was produced in HEK293FT by co-transfection of cDNA containing lentiviral plasmid together with pCMV-dR8.2 dvpr (Addgene, #8455, gift from Bob Weinberg), pCMV-VSV-G (Addgene, #8454, gift from Bob Weinberg) and pAdVAntage (Promega) using FugeneHD (Promega). Supernatants were collected 48h post-transfection, filtered and added to recipient cells in presence of Polybrene (SCBT). Transduced cells were subsequently selected using Puromycin or Hygromycin for 5-7 days.

### Virus propagation and titration

HCoV-OC43 was obtained from ATCC (VR-1558) and propagated in Huh7.5.1 cells at 33C. HCoV-229E was obtained from ATCC (VR-740) and propagated in Huh7.5.1 cells at 33C. SARS-CoV-2 (USA/WA-1/2020 strain) was obtained through BEI Resources (NR-52281) and propagated in Vero cells. Supernatants were collected when cytopathic effect was apparent, filtered and stored at −80C. Viral titers were determined by standard plaque assay using either Huh7.5.1 cells (OC43 and 229E) or Vero cells (SARS-CoV-2). Briefly, serial 10-fold dilutions of virus stocks were used to infect cells in 6-well plates for 1h and an overlay of DMEM media containing 1.2% Avicel RC-591 was added. Cells were incubated for 3-4 days, followed by fixation with 10% formaldehyde, staining with crystal violet and plaque counting. Additionally, SARS-CoV-2 stock was sequence-verified by next-generation sequencing. All experiments with OC43 and 229E were performed in a biosafety level 2 laboratory and all experiments involving SARS-CoV-2 were performed in a biosafety level 3 laboratory.

### Genome-wide CRISPR screens

Huh7.5.1-Cas9 cells were generated by lentiviral transduction with lentiCas9-blast (Addgene, #52962, gift from Feng Zhang) and subsequently selected with blasticidin for 7 days. A portion of Huh7.5.1-Cas9 cells were additionally transduced with lentivirus containing ACE2-IRES-TMPRSS2-hygro. To generate CRISPR KO libraries, a total of 240 million Huh7.5.1-Cas9-blast or Huh7.5.1-Cas9-blast+ACE2-IRES-TMPRSS2-hygro cells were transduced with lentivirus of the human GeCKO v2 library (Addgene, #1000000049, gift from Feng Zhang) at a moi of 0.4 and subsequently selected using puromycin and expanded for 7 days. A total of 60 million mutagenized cells for each GeCKO sublibrary (A and B) were collected for genomic DNA extraction to assess the sgRNA representation of the starting population. For the SARS-CoV-2 CRISPR host factor screen, 100 million cells of Huh7.5.1-Cas9-blast+ACE2-IRES-TMPRSS2-hygro GeCKO library cells were infected with SARS-CoV-2 at a multiplicity of infection (moi) of 0.01. Virus-induced cell death was apparent after 2-3 days and surviving cells were collected 12 dpi. The screen was performed once.

For the 229E and OC43 CRISPR screens, 100 million cells (per screen) of Huh7.5.1-Cas9-blast GeCKO library cells were infected with 229E and OC43 at moi of 0.05 and 3, respectively. Cells were incubated at 33C to increase CPE, which was apparent after 3-4 days. Surviving cells were collected after 10 days for 229E and 14 days for OC43. Each screen was performed in two replicates. For all CRISPR screens, genomic DNA (gDNA) was extracted using either QIAamp DNA Blood Maxi Kit (Qiagen) or Quick-DNA Midiprep Plus (Zymo). The sgRNA expression cassettes were amplified from gDNA in a two-step nested PCR using KAPA HiFi HotStart ReadyMixPCR Kit (Kapa Biosystems). For PCR1, 40 reactions (for control samples) and 10-16 reactions (for virus selected samples) containing 4 μg gDNA were set up and amplified for 16 cycles. Reactions were pooled, mixed and 200 μl were cleaned up using QIAquick PCR Purification kit (Qiagen). For PCR2, 3 reactions containing 5 μl PCR1 product were amplified for 12 cycles using indexed primers. PCR products were gel purified using QIAquick Gel Extraction Kit (Qiagen) and sequenced on an Illumina NextSeq 500 using a custom sequencing primer. Primers sequences are listed in Supplementary Table 4.

Demultiplexed FASTQ files were aligned to a reference table containing sgRNA sequences and abundance of each sgRNA was determined for each starting and selected cell population. Guide count tables were further processed using MaGECK with default “norm-method” to determine positive enrichment scores for each gene ^66^. For 229E and OC43, two biological screen replicates were used as input, and for SARS-CoV-2, one biological screen replicate was used. The gene ontology enrichment of the individual screens was run on genes with MaGECK positive score <= 0.005 using the GO Biological Processes of the Molecular Signatures Database (MSigDB).

### Network propagation

We performed network propagation analysis for the three virus CRISPR screens using the Pathway Commons network ^67^. Specifically, we used a heat-diffusion kernel analogous to random walk with restart (RWR, also known as insulated diffusion and personalized PageRank) which better captures the local topology of the interaction network compared to a general heat diffusion process. The process is captured by the steady-state solution as follows:

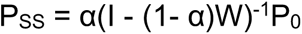

where P_SS_ represents the vector of propagated values at steady-state, P_0_ is the initial labeling (genes of interest from molecular studies), W is the normalized version of the adjacency matrix of the underlying network (in this implementation W = AD^-1^, where A is the unnormalized adjacency matrix, and D is the diagonal degree matrix of the network), I is the identity matrix, and α denotes the restart probability (here, α=0.2), which is the probability of returning to the previously visited node, thus controlling the spread through the network.

We performed three independent propagations, one for each CRISPR dataset (i.e. each virus). After propagation, each propagated network was integrated by multiplying gene-wise. Such an operation is used to create a gene list ranked to prioritize genes with high scores from all propagated datasets. To control for nodes with high degree (i.e. many connections), which due to their heightened connectivity are biased to receive higher propagation scores, we conducted a permutation test. Specifically, we simulated random propagations by shuffling the positive scores to random genes, repeating this 20,000 times per CRISPR screen. Next, we calculated an empirical p-value by calculating the fraction of random propagation runs greater than or equal to the true propagation run for each gene.

The network was created by extracting a subnetwork from the same Pathway Commons network corresponding to genes possessing a significant p-value (p<=0.01) from the propagation (n=378). Of these, interconnected genes were visualized using Cytoscape (n=284). The resulting network was clustered into subnetworks using the GLay Cytoscape plugin ^68^. Three large clusters (1, 3, and 5) were further clustered using GLay into additional subclusters (denoted with letters), resulting in a total of 25 subnetwork clusters (see Extended Data Fig. 3a and Supplementary Table 3). Lastly, Gene Ontology (GO) enrichment analysis (biological process) was performed for each of the 25 resulting subclusters to identify biological processes and pathways associated with each subcluster.

### Generation of clonal Huh7.5.1 KO cell lines

sgRNA sequences against gene targets were designed using the GPP sgRNA Designer (https://portals.broadinstitute.org/gpp/public/analysis-tools/sgrna-design). DNA oligos (IDT) containing sgRNA sequences were annealed and ligated into pX458 (Addgene, #48138, gift from Feng Zhang). Cells were transfected with pX458 constructs using Mirus TransIT-X2 (Mirus Bio) and two days later GFP positive cells were single-cell sorted into 96-well plates using a Sony SH800 cell sorter. For genotyping, genomic DNA was isolated from obtained clones using DNA QuickExtract (Lucigen), the sgRNA-targeted sites PCR amplified and the products Sanger-sequenced. Obtained sequences were compared to reference sequences and clones containing a frameshift indel or de novo stop codon were selected. A list of all used sgRNA sequences and genotyping primers can be found in Supplementary Table 4.

### Generation of RNP edited A549-ACE2 cells

sgRNAs were designed according to Synthego’s multi-guide gene knockout. Briefly, two or three sgRNAs are bioinformatically designed to work in a cooperative manner to generate small, knockout-causing, fragment deletions in early exons. These fragment deletions are larger than standard indels generated from single guides. The genomic repair patterns from a multi-guide approach are highly predictable based on the guide-spacing and design constraints to limit off-targets, resulting in a higher probability protein knockout phenotype.

RNA oligonucleotides were chemically synthesized on Synthego solid-phase synthesis platform, using CPG solid support containing a universal linker. 5-Benzylthio-1H-tetrazole (BTT, 0.25 M solution in acetonitrile) was used for coupling, (3-((Dimethylamino-methylidene)amino)-3H-1,2,4-dithiazole-3-thione (DDTT, 0.1 M solution in pyridine) was used for thiolation, dichloroacetic acid (DCA, 3% solution in toluene) for used for detritylation. Modified sgRNA were chemically synthesized to contain 2’-O-methyl analogs and 3’ phosphorothioate nucleotide interlinkages in the terminal three nucleotides at both 5’ and 3’ ends of the RNA molecule. After synthesis, oligonucleotides were subject to series of deprotection steps, followed by purification by solid phase extraction (SPE). Purified oligonucleotides were analyzed by ESI-MS.

To induce gene knockout, 5 pmol Streptococcus Pyogenes NLS-Sp.Cas9-NLS (SpCas9) nuclease (Aldevron) was combined with 15 pmol total synthetic sgRNA (5 pmol each sgRNA) (Synthego) to form ribonucleoproteins (RNPs) in 20uL total volume with SE Buffer (Lonza). The RNP assembly reaction was mixed by pipetting up and down and incubated at room temperature for 10 minutes.

All cells were dissociated into single cells using TrypLE Express (Gibco), as described above, resuspended in culture media and counted. 100,000 cells per nucleofection reaction were pelleted by centrifugation at 200 x*g* for 5 minutes. Following centrifugation, cells were resuspended in transfection buffer according to cell type and diluted to 2*10^4^ cells/µL. 5 µL of cell solution was added to preformed RNP solution and gently mixed. Nucleofections were performed on a Lonza HT 384-well nucleofector system using program CM-120. Immediately following nucleofection, each reaction was transferred to a tissue-culture treated 96-well plate containing 100µL normal culture media and seeded at a density of 50,000 cells per well.

Two days post-nucleofection, DNA was extracted from using DNA QuickExtract (Lucigen). Amplicons for indel analysis were generated by PCR amplification. PCR products were cleaned-up and analyzed by sanger sequencing. Sanger data files and sgRNA target sequences were input into Inference of CRISPR Edits (ICE) analysis (ice.synthego.com) to determine editing efficiency and to quantify generated indels ^65^. A list of all used sgRNA sequences and genotyping primers can be found in Supplementary Table 4.

### RT-qPCR infection assays

Cells were plated in 96-well plates (in triplicates for each condition) and infected the next day with virus: HCoV-OC43 (moi=3), HCoV-229E (moi=0.05), SARS-CoV-2 (moi=0.01). For infection with HCoVs, cells were harvested 48 hpi, lysates were reverse transcribed and quantitative PCR was performed on a Bio-Rad CFX96 Touch system using the Power SYBR Cells-to-CT kit (Invitrogen) according to the manufacturer’s instructions. 229E and OC43 RNA levels were quantified with virus-specific primer sets and viral RNA levels were normalized to cellular 18S levels.

For SARS-CoV-2 infections, Huh7.5.1 and A549-ACE2 cells were harvested after 24 and 74h, respectively, using 200 μl DNA/RNA Shield (Zymo) to inactivate virus prior to export from the BSL3 laboratory. Samples were extracted using the Quick-DNA/RNA Viral MagBead kit (Zymo) on a Bravo automated liquid handling platform (Agilent). Briefly, the Bravo RNA extraction protocol consists of: 1) 180 μl sample transfer from 2mL deep well to a 1mL deep well plate containing Proteinase K; 2) addition of Zymo Viral DNA/RNA Buffer for sample lysis; 3) Addition of Zymo MagBeads; 4) 10 minute mixing and shaking of samples with lysis buffer and MagBeads; 5) incubation of the mixture on a 96 well ring magnet to collect the beads to a ring at the bottom of the deep well plate; 6) aspiration of the supernatant and dispensing into a 2mL deep well waste plate; 7) addition of wash buffers 1 with mixing; 8) incubation on the 96 well ring magnet; 9) aspiration. Steps 7-9 are repeated for wash buffer 2 and two rounds of 100% ethanol. 10) incubation on the magnet for 20 minutes to fully evaporate residual 100% ethanol from the beads; 11) Elution with nuclease-free water.

For RT-qPCR, separate reactions were performed for the quantification of SARS-CoV-2 N and E gene transcripts as well as cellular RNaseP for normalization using the Luna Universal Probe One-Step RT-qPCR Kit (NEB) on a Bio-Rad CFX384 Touch system. N and E gene transcripts showed high concordance and N gene levels normalized to RNaseP were displayed in figures. All qPCR primer/probe sequences are listed in Supplementary Table 4.

### Western blots

Cells were lysed using Laemmli SDS sample buffer containing 5% beta-mercaptoethanol and boiled at 95C for 10min. Lysates were separated by SDS-PAGE on pre-cast Bio-Rad 4-15% poly-acrylamide gels in Bio-Rad Mini-Protean electrophoresis system. Proteins were transferred onto PVDF membranes using Bio-Rad Trans-Blot Turbo transfer system. PVDF membranes were blocked with PBS buffer containing 0.1% Tween-20 and 5% non-fat milk. Blocked membranes were incubated with primary antibody diluted in blocking buffer and incubated overnight at 4C on a shaker. Primary antibodies were detected by incubating membranes with 1:5000 dilution of HRP-conjugated (Southern Biotech) secondary anti-mouse and anti-rabbit antibodies for 1 h at room temperature. Blots were visualized using a ChemiDoc MP Imaging System (Bio-Rad). The following primary antibodies and their dilutions were used in this study: GAPDH (SCBT, sc-32233) at 1:1000, ACE2 (R&D Systems, AF933) at 1:1000, TMPRSS2 (Abcam, ab92323) at 1:1000.

### Pseudo-typed virus infection

Cells were plated in 96-well plates and infected with 30 μl of SARS-CoV-2 Reporter Virus Particles (Integral Molecular, RVP-701) per well. After 48-72h, infection rates were measured according the GFP levels using a Cytoflex flow cytometer (Beckman Coulter Life Sciences).

### Compounds

The following compounds were used in this study: SAR405 (SelleckChem, S7682), YM201636 (SelleckChem, S1219), PF-429242 dihydrochloride (Sigma, SML0667) and Fatostatin HBr (SelleckChem, S8284). All compounds were resuspended in DMSO and stored at −20C until use.

### Cell viability assay

Huh7.5.1 cells were treated with compounds at the same concentrations and durations as in infection assays. Cell viability was measured using Cell Titer Glo (Promega) according to the manufacturer’s instructions.

## Supplementary Tables

Table 1: CRISPR screen results. MaGECK output for positive gene enrichment analysis of SARS-CoV-2, 229E and OC43 host factors.

Table 2: Gene ontology enrichment analysis of individual CRISPR screens and network propagation clusters.

Table 3: Network propagation results.

Table 4: DNA oligo sequences used in this study.

## Data availability

Raw sequencing data for CRISPR KO screens will be made available through the EMBL-EBI ArrayExpress (https://www.ebi.ac.uk/arrayexpress/).

## Acknowledgments

This research was funded by grants from the National Institutes of Health (P50AI150476, U19AI135990, U19AI135972, R01AI143292, R01AI120694, P01A1063302, and R01AI122747 to N.J.K.; F32CA239333 to M.B.; 5DP1DA038043 to M.O.). M.O. acknowledges support through a gift from the Roddenbury. Funding to A.S.P. was provided by the Chan Zuckerberg Biohub. We would like to thank Dr. Anita Sil, Dr. Bastian Joehnk, Dr. Lauren Rodriguez, and Keith Walcott for BSL-3 laboratory support; Matthew Laurie for help with Bravo liquid handling; Dr. Sandra Schmid, Dr. Don Ganem and Dr. Francoise Chanut for editorial comments; Dr. Olga Gulyaeva, Sara Sunshine, Dr. Marco Hein, Dr. Scott Biering, Dr. Nathan Meyers, and Dr. Jan Carette for helpful discussion. We also express gratitude to the Biohub lab management and operations team for their support.

## Author contributions

R.W., C.R.S., J.K., K.T., J.M.H., J.C.S., J.O., and A.S.P. performed the experiments. R.W., C.R.S., J.K. and A.S.P. designed experiments. R.W., J.K., M.B. and A.S.P. analyzed and visualized data. N.J.K., M.O., K.H. and A.S.P. supervised study and provided technical guidance. A.S.P. conceptualized study and wrote initial draft of the manuscript. All authors provided comments and edits on the manuscript.

